# Integrated analysis of spatial multi-omics with SpatialGlue

**DOI:** 10.1101/2023.04.26.538404

**Authors:** Yahui Long, Kok Siong Ang, Sha Liao, Raman Sethi, Yang Heng, Chengwei Zhong, Hang Xu, Nazihah Husna, Min Jian, Lai Guan Ng, Ao Chen, Nicholas RJ Gascoigne, Xun Xu, Jinmiao Chen

## Abstract

Integration of multiple data modalities in a spatially informed manner remains an unmet need for exploiting spatial multi-omics data. We introduce SpatialGlue, a graph neural network with dual-attention mechanism, to learn each modality’s significance at cross-omics and intra-omics integration. We demonstrate that SpatialGlue can accurately aggregate cell types into spatial domains at a higher resolution on different tissue types and technology platforms, as well as gain insights into cross-modality spatial correlations.

## Main

Spatial transcriptomics is the next major development in analyzing biological samples since the advent of single-cell transcriptomics. Currently, spatial technologies are expanding to spatial multi-omics with simultaneous profiling of different omics on a single tissue section. These technologies can be roughly divided into two categories, sequencing-based and imaging-based. Sequencing-based techniques include DBiT-seq ^1^, spatial-CITE-seq ^2^, spatial-ATAC-RNA-seq and CUT&Tag-RNA-seq ^3^, SPOTS ^4^, SM-Omics ^5^, Stereo-CITE-seq ^6^, and spatial RNA-TCR-seq ^7^ while imaging-based techniques include DNA seqFISH+ ^8^, and DNA-MERFISH based DNA and RNA profiling ^9^. To fully utilize spatial multi-omics data and construct a coherent picture of the tissue under study, spatially aware integration of heterogeneous data modalities is required. Multi-omics data integration poses a significant challenge as different modalities have feature counts that can vary enormously (e.g., protein vs transcripts) and possess different statistical distributions. This challenge is deepened when integrating spatial information with feature counts within each data modality. To our knowledge, there is no tool designed specifically for spatial multi-omics. Existing tools such as SpaGCN ^10^ and GraphST ^11^ target spatial single omics integrated analysis, while GLUE ^12^ and Seurat WNN ^13^ perform multi-omics data integration without employing spatial information.

Here we introduce SpatialGlue, a graph neural network (GNN) based deep learning model that performs spatial multi-omics data integration (Figure 1a). SpatialGlue employs attention aggregation to achieve data integration on two levels, within-modality spatial information and measurement feature integration, and between-modality integration. SpatialGlue first learns a low dimension embedding space within each modality using spatial and omics data. Within each modality, SpatialGlue constructs a spatial proximity graph and a feature graph which are used separately to encode the pre-processed expression data into a common low dimension embedding space. Here the spatial proximity graph captures spatial relationships between measurement spots, while the feature graph captures feature similarities between spots that can be spatially distant. These constructed graphs can possess unique semantic information that should be integrated. Therefore, we adopted a within-modality attention aggregation layer to adaptively integrate the spatial and feature graph-specific representations and derive modality-specific representations. Specifically, the model learns graph-specific weights to assign importance to each graph. Similarly, the different data modalities can have distinct and complementary contributions to each spot. Thus, we further designed a between-modality attention aggregation layer that learns modality-specific importance weights and adaptively integrates the modality-specific representations to generate the final cross-modality integrated latent representation. The learned weights illustrate the contribution of each modality to the learned latent representation of each spot and consequently the demarcation of different cell types. We believe this approach enables more accurate integration than the common approaches of summation or concatenation.

**Figure 1:**
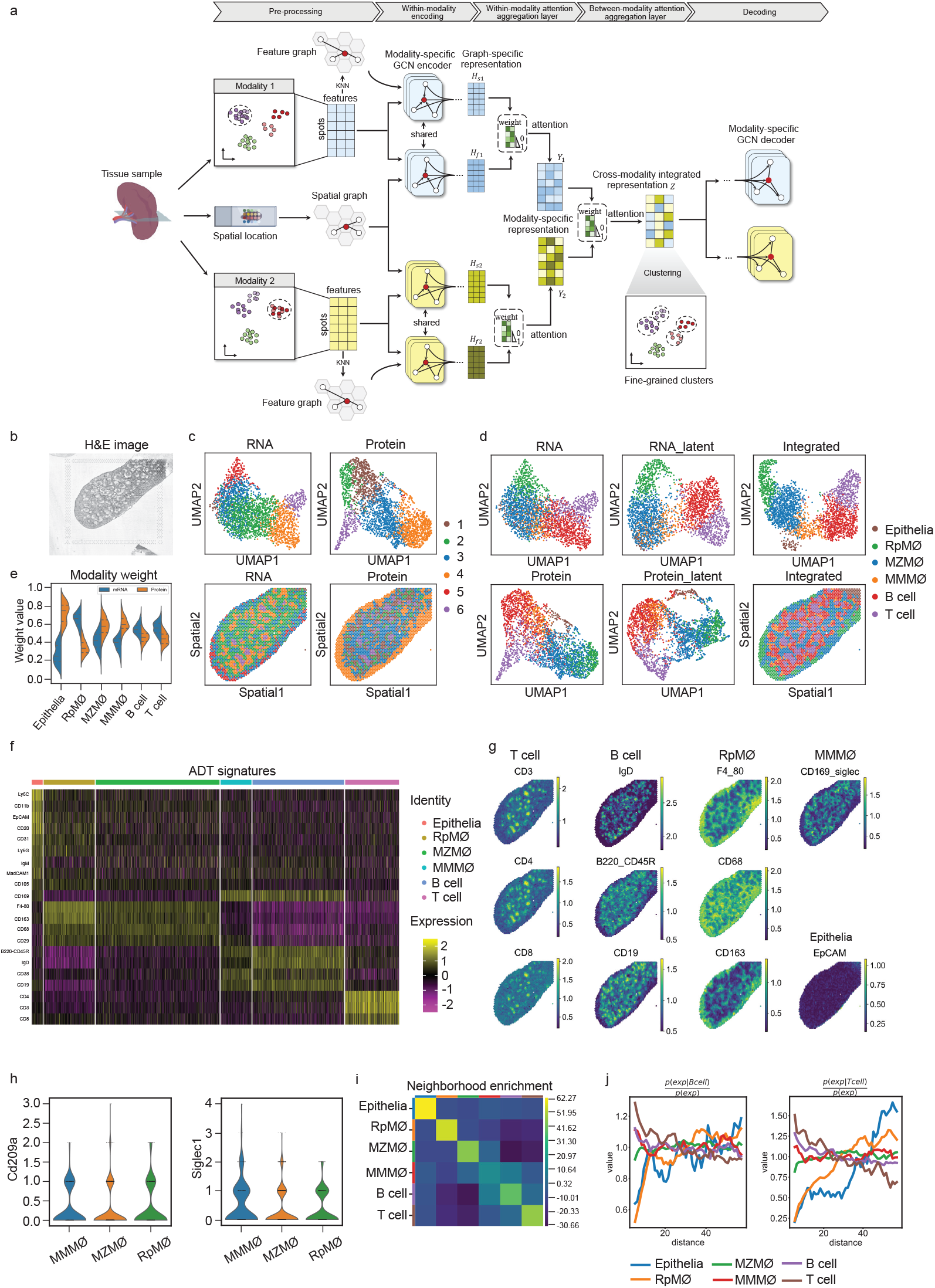
Interpretable deep dual-attention model enables the identification of fine-grained cell types in mouse spleen data generated using SPOTS. (a) Overview of the SpatialGlue framework. A spatial multi-omics technology simultaneously measures two distinct types of molecules, e.g., RNA and surface protein, while preserving spatial context of the tissue. SpatialGlue first uses the *K*-nearest neighbor (KNN) algorithm to construct a spatial neighbor graph using the spatial coordinates and a feature neighbor graph with the normalized expression data for each omics modality. Each modality has an implemented GNN-encoder that takes its normalized expressions and neighbor graph to learn two graph-specific representations by iteratively aggregating representations of neighbors. To capture the importance of different graphs, we designed a Within-Modality attention aggregation layer to adaptively integrate graph-specific representations and obtain a modality-specific representation. Finally, to preserve the importance of different modalities, SpatialGlue uses a Between-Modality attention aggregation layer to adaptively integrated modality-specific representations and output the final integrated representation of spots. (b) H&E image of the mouse spleen. (c) UMAP plots and spatial clustering of the RNA and protein expression data. (d) Comparison of UMAP plots of RNA and protein modalities and SpatialGlue integrated output, all colored by annotated clusters obtained from the integrated output. (e) Modality weights explaining the importance of different modalities to each cluster in the mouse spleen dataset. (f) Heatmap showing the expression levels of differentially expressed ADTs for each cluster. (g) Normalized ADT levels of key surface markers for T cell (CD3, CD4, CD8), B cell (IgD, B220, CD19), RpMΦ (F4_80, CD68, CD163), MMMΦ (CD169), and Epithelia (EpCAM). (h) Violin plots indicating the expression distribution of two marker genes in the MMMΦ, MZMΦ, and RpMΦ clusters. (i) Neighborhood enrichment of cell type pairs. (j) Cluster co-occurrence score for each cluster at increasing distances. The full names of the abbreviations RpMΦ, MMMΦ, and MZMΦ are red pulp macro, CD169^+^ MMM, CD209a^+^ MZM, respectively.

We first demonstrate SpatialGlue’s capabilities with murine spleen spatial profiling data consisting of protein and transcript measurements ^4^. The spleen is an important organ within the lymphatic system with functions including B cell maturation in germinal centers formed within B cell follicles. These are complex structures with an array of immune cells present (Figure 1b). The data was generated using SPOTS with the 10x Visium technology capturing whole transcriptomes and extracellular proteins via polyadenylated antibody-derived tag-conjugated (ADT-conjugated) antibodies. The protein detection panel was designed to detect the surface markers of B cells, T cells, and macrophages which are well represented in the spleen. After preprocessing, we performed clustering of each data modality and plotted the clusters on the tissue slide to examine their correspondence between modalities (Figure 1c). The clusters clearly did not align, indicating that each modality possessed different information content. We next integrated both modalities with the spatial context, followed by clustering (Figure 1d and Supplementary Figure S1a). Using the protein markers and DEGs, clusters of spots enriched with B cells, T cells, macrophage subsets, and epithelia were annotated ^14–16^ (Figure 1f-h and Supplementary S1b). In particular, we could identify epithelial cells and macrophage subsets that were not annotated in the original study. We also plotted the clusters obtained from the integrated analysis onto the individual modalities’ UMAPs to examine their respective contributions. The epithelia enriched spots were better separated in the protein modality but were mixed with marginal zone macrophage (MZMΦ) enriched spots in the RNA modality. This contribution to separation by the protein modality can be seen in the learned modality weights (Figure 1e). Conversely, the red pulp macrophage (RpMΦ) enriched spots were better separated in the RNA UMAP and likewise showed greater contribution in the RNA modality weight. For the remaining cell type enriched spots such as marginal metallophilic macrophages (MMMΦ), the spots were more mixed in the individual modalities and better separated in the integrated analysis. Specifically, the MZMΦ spots were mixed with MMMΦ in the RNA modality and RpMΦ in the protein modality. By leveraging on both modalities, our model was able to separate the different cell types.

Within the white pulp zone, the T cell spots were concentrated in small clusters known as T cell zones. The B cell enriched spots were mainly found in areas adjacent to the T cell clusters. We also visualized the spatial distribution of the cell types’ protein markers (Figure 1g). For the B cells, their signal (CD19) indicated that they were much more widely distributed within the spleen with naïve B cells (IgD^+^) colocalized in clusters surrounding the T cell zones. The RpMΦ markers were unsurprisingly the strongest in the red pulp zone while the MMMΦ marker (CD169/Siglec) was expressed in areas surrounding the B and T cell zones. Expectedly, the RpMΦ and MMMΦ marker expressions were mostly mutually exclusive spatially. Finally, EpCAM (epithelia) expression was only found in only a small portion of the slide at the top right where the epithelia spots were annotated. We also examined the markers, Cd209a (MZMΦ) and Siglec1 (MMMΦ) in the RNA modality for their expression by the macrophage clusters, confirming their identity (Figure 1h).

From the cluster and marker visualization, we observed cell types whose zones were adjoining. Thus, we quantitated the spatial relationship by computing neighborhood enrichment (Figure 1i) and co-occurrence scores with respect to distance from the T and B cell perspective (Figure 1j). In general, we observed a neighborhood enrichment among the B cells, T cells, and MMMΦ. The co-occurrence scores also showed similar trends with B and T cell spots being most likely to be found together at the closest distance. This was followed by MMMΦ which surrounded T and B cell clusters in the white zone. These reflect the layers of cell types that form the follicles and their surroundings. We also note a neighborhood enrichment of MZMΦ and MMMΦ due to the MZMΦ being positioned within the marginal zone surrounding white pulp which in turn was enriched with MMMΦ. We next compared SpatialGlue’s output with Seurat WNN’s (Supplementary Figure S1c). Compared to SpatialGlue, Seurat WNN assigned a continuous border zone of RpMΦ (cluster 3). Examining the RpMΦ markers F4/80 and CD163, they did not correspond to a similar continuous zone (Figure 1g). Instead, these markers showed a discontinuous and less concentrated boundary layer. Within the white pulp, the T cell zone (red spots) was easily distinguished but the B cell and MMMΦ consecutive layering was less distinct compared to SpatialGlue’s clusters. In terms of modality weights, Seurat WNN assigned higher weights for all clusters to the protein modality (Supplementary Figure S1d), in contrast to SpatialGlue.

We next tested SpatialGlue on a mouse thymus dataset acquired with Stereo-CITE-seq spatial multi-omics ^6^ that measures mRNA and protein at sub-cellular resolution. The thymus is a small gland surrounded by a capsule of fibers and collagen (Figure 2a, b). It is divided into two lobes connected by a connective isthmus with each lobe being broadly divided into a central medulla surrounded by an outer cortex layer. In each data modality, broad outlines of the medulla regions and the surrounding cortex could be seen (Figure 2c). However, clusters in the RNA modality did not capture the cortex layers while the clusters in the protein modality were more fragmented. The integrated results showed much more contiguous clusters which we could easily annotate (Figure 2d and Supplementary Figure S2a). Here the cortex was divided into three regions, inner (3), middle (4), and outer (5), and was separated from the medulla by the corticomedullary junction (CMJ) ^17^. We then confirmed the annotation with the markers and DEGs of cell types expected in each region (Supplementary Figure S2b, c). For most regions, the RNA modality made the greatest contributions, except for the middle cortex (4) and subcapsular zone (7) (Figure 2e). The protein modality’s contribution to distinguishing the middle cortex is visible in Figure 2c with the corresponding cluster 3. We again compared SpatialGlue to Seurat WNN on data integration. For this dataset, the Seurat WNN output captured the major regions in the thymus, but the clusters showed greater fragmentation, especially in the cortex (Supplementary Figure S2d). Like the spleen example, Seurat WNN also assigned higher modality weights to the protein modality for all clusters that it found (Supplementary Figure S2e).

**Figure 2:**
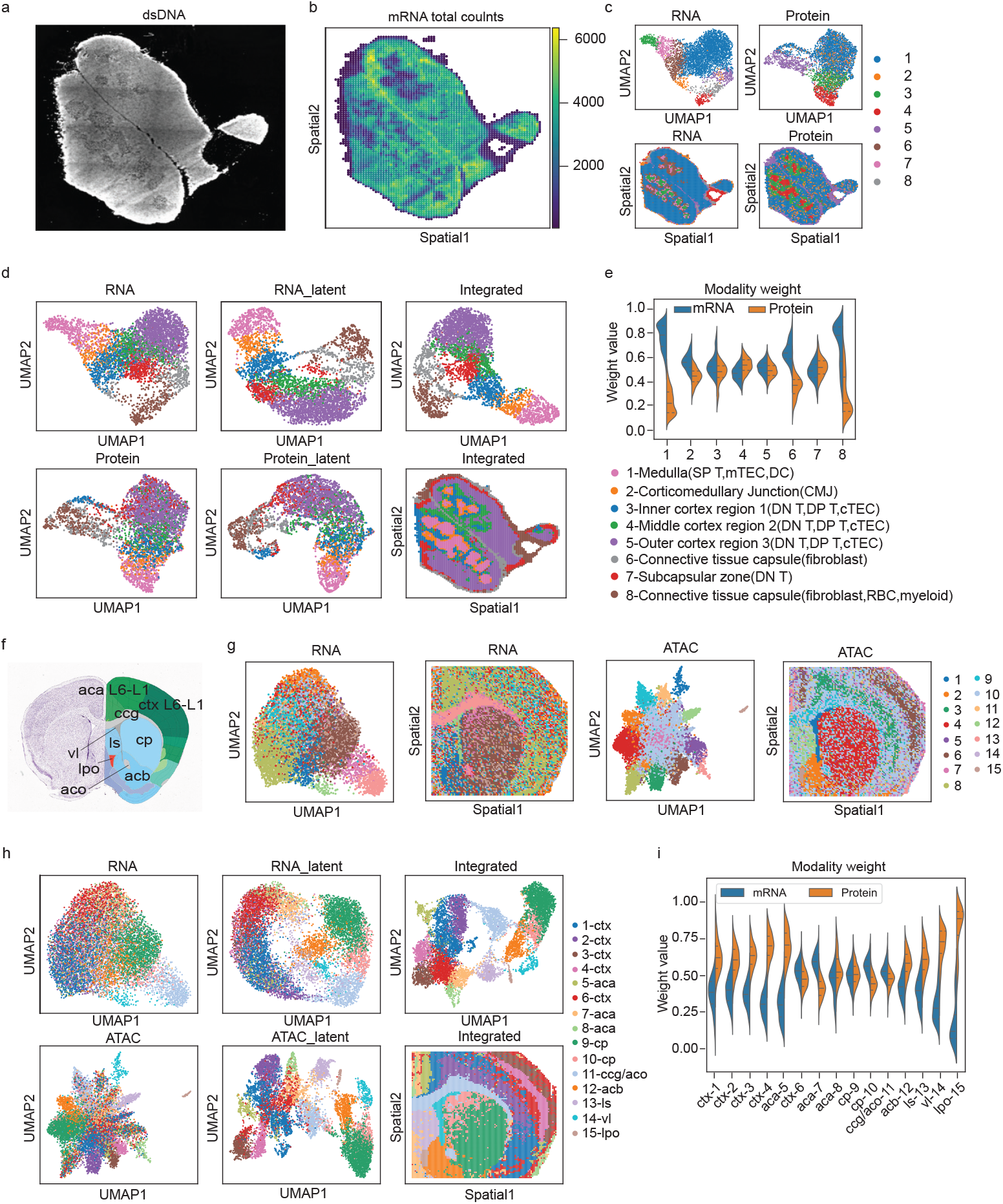
SpatialGlue accurately integrates different datasets of mouse thymus and mouse brain. The mouse thymus data of protein and RNA modalities was acquired with Stereo-CITE-seq, and the mouse brain data of RNA and ATAC modalities was acquired with spatial-ATAC-RNA-seq. (a) dsDNA image of the mouse thymus. (b) Total mRNA counts in the mouse thymus dataset. (c) UMAP plots and spatial clustering of the RNA and protein expression data in the mouse thymus dataset. (d) Comparison of UMAP plots of RNA and protein modalities and SpatialGlue integrated output in the mouse thymus dataset, all colored by annotated clusters obtained from the integrated output. (e) Modality weights explaining the importance of each modality to each cluster in the mouse thymus dataset. (f) Annotated reference mouse brain coronal section from Allen Mouse Brain Atlas. (g) UMAP plots and spatial clustering of the RNA expression and ATAC data in the mouse brain dataset. (h) Comparison of UMAP plots of RNA and ATAC modalities and SpatialGlue integrated output in the mouse brain dataset, all colored by annotated clusters obtained from the integrated output. (i) Modality weights explaining the importance of each modality to each cluster in the mouse brain dataset. The full names of abbreviation used in the annotations of (f) and (h) are, ctx: cerebral cortex, cp: caudoputamen, vl: lateral ventricle, lpo: lateral preopic area, aca: anterior cingulate area, ls: lateral septal nucleus, aco: anterior commissure, olfactory limb, acb: nucleus accumbens, cc: corpus callosum.

In our final example, we tested SpatialGlue on a P22 mouse brain coronal section dataset acquired using spatial ATAC-RNA-seq ^3^ to measure mRNA and open chromatin regions. We employed the Allen brain atlas reference to annotate anatomical regions such as the cortex layers (ctx), genu of corpus callosum (ccg), lateral septal nucleus (ls), and nucleus accumbens (acb) (Figure 2f). Analyzing individual modalities, we see that they captured various regions with differing accuracy. While both modalities captured the caudoputamen (cp), lateral ventricle (vl), and olfactory limb of the anterior commissure (aco), the RNA modality captured the ccg but was unable to differentiate the ctx layers or the ls (Figure 2g). Meanwhile, the ATAC modality was able to isolate the ls and even the lateral preoptic area (lpo), as well as portions of the acb and some of the ctx layers. With SpatialGlue, the integrated analysis captured all the aforementioned anatomical regions and produced better defined layers in the ctx and anterior cingulate area (aca) regions (Figure 2h and Supplementary Figure S3a). We also found DEGs in the expected brain regions, such as Cux2 and Olfm1 in the cortex layers (ctx-3,4), and myelin related genes, Tspan2, Cldn11 and Ugt8a expressed in the post-natal developing corpus callosum (11-ccg/aco) (Supplementary Figure 3b). The contributions by each modality to the integrated result for each cluster were illustrated in the learned modality weights (Figure 2i). As expected from the individual analysis, the ATAC modality was the key contributor towards distinguishing many of the ctx layers (1, 2, 3, 4), acb (12), ls (13), and lpo (15). For the RNA modality, it made a greater contribution to the ccg/aco (11), aca (7), and cp (10) clusters. Again, these results demonstrated SpatialGlue’s ability to extract the relevant information from each modality and integrate them in a spatially aware manner to capture anatomical regions with superior accuracy. Lastly, we compared SpatialGlue to Seurat WNN. Seurat WNN’s output also captured many anatomical regions like the ctx layers, ccg, vl, and cp, but did not differentiate the lpo from the acb (Supplementary Figure S3c). Moreover, the Seurat WNN output was grainy without clear boundaries between regions. Seurat WNN also heavily relied on the ATAC data in the data integration but to a much greater extent than SpatialGlue (Supplementary Figure S3d).

With these three examples, we demonstrated SpatialGlue’s ability to effectively integrate multiple data modalities within their spatial context to reveal histologically relevant structures of tissue samples. Our examples spanned different tissue types and different acquisition technologies, which highlighted its applicability to a wide range of data. SpatialGlue was designed to be computation resource efficient. The largest dataset we tested contained 9,215 spots (spatial-ATAC-RNA-seq mouse brain), and it required about 5 mins of wall-clock time on a server with Intel Core i7-8665U CPU and NVIDIA RTX A6000 GPU. Therefore, we believe SpatialGlue will be an invaluable analysis tool for present and future spatial multi-omics data. Although the examples so far only include two omics data modalities, the framework is extensible to three or more. As data with greater numbers of modalities become available, we will demonstrate SpatialGlue’s applicability. We also plan to extend SpatialGlue’s functionality with integration of multi-omics data acquired from adjacent tissue slices.

## Methods

### Data

*SPOTS mouse spleen dataset* Ben-Chetrit et al. ^4^ processed fresh frozen mouse spleen tissue samples and analyzed them using the 10x Visium system supplemented with DNA-barcoded antibody staining. The antibodies (poly(adenylated) antibody-derived tags (ADTs)) enabled protein measurement alongside the transcriptome profiling by 10x Visium. The panel of 21 ADTs was designed to capture the markers of immune cells found in the spleen, including B cells, T cells, and macrophages. A total of 2,653 spots were captured with 32,285 genes per spot. In this example, we used replicate one. We first filtered out genes expressed in fewer than 10 cells. The filtered gene expression counts were then log-transformed and normalized by library size using the SCANPY package. Finally, the top 3,000 HVGs were selected and used as input for PCA. We used the first 50 principal components as the input of the encoder to ensure a consistent input dimension with the ADT data. For the ADT data, we applied CLR normalization to the raw protein expression counts. PCA was then performed on the normalized data and the top 50 principal components were used as input to the encoder.

*Stereo-CITE-seq mouse thymus dataset* Murine thymus tissue sample was investigated with the Stereo-CITE-seq spatial multi-omics by Liao et al. ^6^. The acquired data consisted of 4,697 spots with 23,622 genes and 51 proteins. For the transcriptomic data, we first filtered out genes expressed in fewer than 10 cells and spots with fewer than 80 gene expressed. The filtered gene expression counts were next log-transformed and normalized by library size via the SCANPY package ^18^. Finally, to reduce the dimensionality of the data, the top 3,000 highly variable genes (HVGs) were selected and used as input for PCA. To ensure a consistent input dimension with the ADT data, the first 22 principal components were retained and used as the input of the encoder. For the ADT data, we first filter out proteins expressed in fewer than 50 cells, resulting 22 proteins retained. The protein expression counts were then normalized using CLR (Centered Log Ratio) across each cell. PCA was then performed on the normalized data, and all 22 principal components were used as the input of the encoder.

*Spatial-ATAC-RNA-seq mouse brain dataset* This example utilized a juvenile (P22) murine brain tissue sample analyzed by Zhang et al. with Spatial-ATAC-RNA-seq ^3^. Microfluidic barcoding was used to capture spatial location and combined with in situ Tn5 transposition chemistry to capture chromatin accessibility. The data captured consisted of 9,215 spots with 2,2914 genes and 1,210 peaks. To preprocess the transcriptomic data, cells expressing fewer than 200 genes and genes expressed fewer than 200 cells were filtered out. Next, the gene expression counts were log-transformed and normalized by library size via the SCANPY package. The top 3,000 highly variable genes (HVGs) were selected and used as input to PCA for dimensionality reduction. For consistency with the chromatin peak data, the first 50 principal components were retained and used as input to the encoder. For the chromatin peak data, we used LSI (latent semantic indexing) to reduce the raw chromatin peak counts data to 50 dimensions.

### The SpatialGlue framework

We first consider a spatial multi-omics dataset with two different omics modalities, each with a distinct feature set 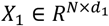 and 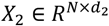. *N* denotes the number of spots in the tissue. *d*_1_ and *d*_2_ are the numbers of features for two omics modalities, respectively. For example, in spatial-ATAC-RNA-seq, *X*_1_ and *X*_2_ refer to the sets of genes and chromatin regions respectively, while in Stereo-CITE-seq, *X*_1_ and *X*_2_ refer to the sets of genes and proteins respectively. The primary objective of spatial multi-omics data integration is to learn a mapping function that can project the original individual modality data into a uniform latent space and then integrate the resulting representations. As shown in Figure 1a, the SpatialGlue framework consists of four major modules: (1) Modality-specific GCN encoder, (2) Within-Modality attention aggregation layer, (3) Between-Modality attention aggregation layer, and (4) Modality-specific GCN decoder. The details of each module are described next. Notably, here we demonstrate the SpatialGlue framework with two modalities. Benefiting from the modular design, SpatialGlue readily extends to spatial multi-omics data with more than two modalities.

### Construction of neighbor graph

Assuming spots that are spatially adjacent in a tissue usually have similar cell types or cell states, we convert the spatial information to an undirected neighbor graph *G*_*s*_ = (*V, E*) with *V* denoting the set of *N* spots and *E* denoting the set of connected edges between spots. Let *A*_*s*_ ∈ *R*^*N*×*N*^ be the adjacent matrix of graph *G*_*s*_, where *A*_*s*_(*i, j*) = 1 if and only if the Euclidean distance between spots *i* and *j* is less than specific neighbor number *l*, otherwise 0. In our examples, we select the top *r* = 6 nearest spots as neighbors of a given spot for all datasets.

In a complex tissue sample, it is possible for spots with the same cell types to be spatially non-adjacent to each other, or even far away. To capture the proximity of such spots in a latent space, we explicitly model the relationship between them using a feature graph. Specifically, we apply the *k*-nearest neighbor algorithm (KNN) on the PCA embeddings and construct the feature graph 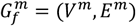, where *V*^*m*^ and *E*^*m*^ denote the sets of *N* spots and connected edges between spots in the *m* ∈ {1,2}-th modality, respectively. For a given spot, we choose the top *k* nearest spots as its neighbors. By default, we set *k* to 20 for all datasets. We use 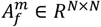 to denote the adjacency matrix of the feature graph 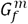. If spot *j* ∈ *V*^*m*^ is the neighbor of spot *i* ∈ *V*^*m*^, then 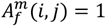, otherwise 0.

### Graph convolutional encoder for individual modality

Each modality (e.g., mRNA or protein) contains a unique feature distribution. To encode each modality in a low dimension embedding space, we use the graph convolution network (GCN) ^19^, an unsupervised deep graph network, as the encoder of our framework. The main advantage of GCNs is that it can capture the cell expression patterns and neighborhood microenvironment while preserving the high-level global patterns. For each modality, using the pre-processed features as inputs, we separately implement a GCN-encoder on the spatial adjacency graph *G*_*s*_ and the feature graph *G*_*f*_ to learn graph-specific representations. These two neighbor graphs reflect distinct topological semantic relationships between spots, enabling the encoder to capture different local patterns and dependencies of each spot by iteratively aggregating the representations from its neighbors. Specifically, the *l* -th (*l* ∈ {1,2, …, *L* − 1, *L*}) layer representation in the encoder are formulated as follows:

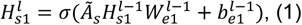

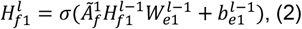

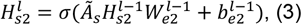

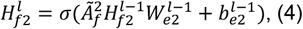

Where 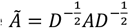 represents the normalized adjacency matrix of specific graph and *D* is a diagonal matrix with diagonal elements being 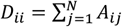. In particular, Ã_*s*_, 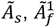, and 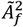 are the corresponding normalized adjacency matrices of the spatial graph, the feature graph 1, and the feature graph 2, respectively. *W*_*e*._, and *b*_*e*._ denote a trainable weight matrix and a bias vector, respectively. *σ*(·) is a nonlinear activation function such as ReLU (Rectified Linear Unit). 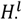 denotes the *l* -th layer output representation, and 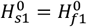 and 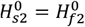 are set as the original input PCA embedding *X*_1_ and *X*_2_ respectively. We also specify 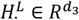, the output at the *L*-th layer, as the final latent representation of the encoder with *d*_3_ as the hidden dimension. *H*_*sk*_ and *H*_*fk*_ represent the latent representations derived from the spatial and feature graphs within modality k, respectively.

### Within-Modality attention aggregation layer

For an individual modality, taking its pre-processed features and two graphs as input, we can derive two graph-specific spot representations via the graph convolutional encoder, such as *H*_*s*1_ and *H*_*f*1_. To integrate the graph-specific representations, we design a Within-Modality attention aggregation layer following the encoder such that its output representation preserves expression similarity and spatial proximity. Given that different neighbor graphs can provide unique semantic information for each spot, the aggregation layer is designed to integrate graph-specific representations in an adaptive manner by capturing the importance of each graph. As a result, we derive a modality-specific representation for each modality. Specifically, for a given spot *i*, we first subject its graph-specific representation 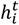 to a linear transformation (i.e., a fully connected neural network), and then evaluate the importance of each graph by the similarity of the transformed representation and a trainable weight vector *q*. Formally, the attention coefficient 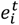, representing the importance of graph *t* to the spot *i*, is calculated by:

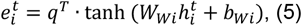

where *W*_*Wi*_ and *b*_*Wi*_ are the trainable weight matrix and bias vector, respectively. To reduce the number of parameters in the model, all the trainable parameters are shared by the different graph-specific representations. To make the attention coefficient comparable across different graphs, a softmax function is applied to the attention coefficient to derive attention score 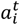.

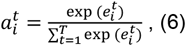

where *T* denotes the number of neighbor graphs (set to 2). 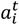 represents the semantic contribution of the *t*-th neighbor graph to the representation of spot *i*. A higher value of 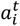 means greater importance.

Subsequently, the final representation 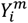 of spot *i* in the *m* -th modality can be generated by aggregating graph-specific representations according to their attention scores:

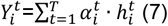

such that 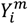 preserves the raw cell expressions, cell expression similarity, and spatial proximity within modality *m*.

### Between-Modality attention aggregation layer

Each individual omic modality provides a partial view of a complex tissue sample, thus requiring an integrated analysis to obtain a comprehensive picture. These views can contain both complementary and contradictory elements, and thus different importance should be assigned to each modality to achieve coherent data integration. Here we use a Between-Modality attention aggregation layer to adaptively integrate the different data modalities. This attention aggregation layer will focus on the more important omics modality by assigning greater weight values to the corresponding representation. Like the Within-Modality layer, we first learn the importance of modality *m* by calculating the following coefficient 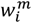:

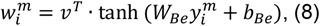

Where 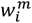 are attention coefficients that represent the importance of modality *m* to the representation of spot *i. W*_*Be*_, *b*_*Be*_, and *v* are learnable weight and bias variables, respectively. Similarly, we further normalize the attention coefficients using the softmax function:

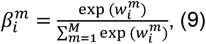

where 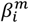 is the normalized attention score that represents the contribution of modality *m* to the representation of spot *i*.

Finally, we derive the final representation *Z*_*i*_ of spot *i* by aggregating each modality-specific representation according to their attention score 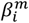:

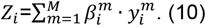

After model training, the latent representation 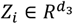 can be used in various downstream analyses, including clustering, visualization, and DEG detection.

### Model training

The resulting model is trained jointly with three different loss functions for reconstruction loss, correspondence loss, and adversarial loss. Each loss function is described as follows.

*Reconstruction loss* To enforce the learned latent representation to preserve the expression profiles from different modalities, we design an individual decoder for each modality to reverse the integrated representation *Z*_*i*_ back into the normalized expression space. Specifically, by taking output *Z*_*i*_ from the Between-Modality attention aggregation layer as input, the reconstructed representations 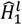 and 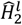 from the decoder at the *l*-th (*l* ∈ {1,2, …, *L* − 1, *L*}) layer are formulated as follows:

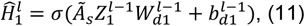

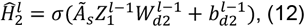

where *W*_*d*1_, *W*_*d*2_, *b*_*d*1_, and *b*_*d*2_ are trainable weight matrices and bias vectors, respectively. 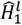 and 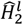 represent the reconstructed expression matrices for omics modalities 1 and 2, respectively.

SpatialGlue’s objective function to minimize the expression reconstruction loss is as follows:

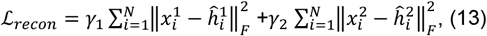

where *x*^1^ and *x*^2^ represent the original features of modalities 1 and 2, respectively. *γ*_1_ and *γ*_2_ are weight factors that are utilized to balance the contribution of different modalities. Due to the differences of sequencing technologies and molecular types, the feature distributions of different omics assays can vary significantly. As such, the weight factors also vary between different spatial multi-omics technologies but are fixed for datasets obtained using the same omics technology.

*Correspondence loss* While reconstruction loss can enforce the learned latent representation to simultaneously capture the expression information of different modality data, it does not guarantee that the representation manifolds are fully aligned across modalities. To deal with the issue, we add a correspondence loss function. Correspondence loss aims to force consistency between a modality-specific representation *Y*_*m*_ and its corresponding representation *Ŷ*_*m*_ obtained through the decoder-encoder of another modality. Mathematically, the correspondence loss is defined as follows:

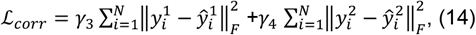

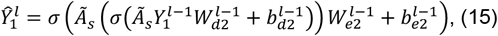

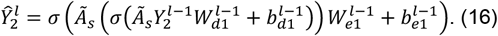

where *γ*_3_ and *γ*_4_ are hyper-parameters, controlling the influences of different modality data. We set 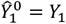 and 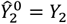. *σ*(·) is a nonlinear activation function, i.e., ReLU (Rectified Linear Unit).

*Adversarial loss* To constrain the representations across different modalities in a uniform latent space, we further add a discriminator, i.e., fully connected neural network, in our model. This discriminator tries to align the representations with *δ* = 1 denoting modality 1 and *δ* = 0 denoting modality 2. Formally, the adversarial loss of the discriminator is defined as:

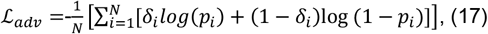

where *p*_*i*_ and (1 − *p*_*i*_) represents the probability scores of the representations assigned to modalities 1 and 2, respectively.

Therefore, the overall loss function used for model training is defined as:

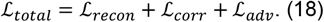

### Implementation details

For all datasets, a learning rate of 0.0001 was used. To account for differences in feature distribution across the datasets, a tailored group of weight factors [*γ*_1_, *γ*_2_, *γ*_3_, *γ*_4_] was assigned to each one. The weight factors were [1, 50, 1, 5] for the SPOTS mouse spleen dataset, [1, 10, 1, 10] for the Stereo-CITE-seq mouse thymus dataset, [1, 2.5, 1, 1] for the spatial-ATAC-RNA-seq mouse brain dataset. The training epochs used for the SPOTS mouse spleen, Stereo-CITE-seq mouse thymus, and spatial-ATAC-RNA-seq mouse brain datasets were 900, 1500, and 1500, respectively.

### Seurat WNN Analysis

For the SPOTS mouse spleen and Stereo-CITE-seq mouse thymus datasets, we employed the same data preprocessing steps as SpatialGlue and used the same PCA coefficients as input to analysis with Seurat WNN (FindMultiModalNeighbors function) ^13^. We then adjusted the resolution input to the Seurat FindCluster function to obtain the same number of clusters as SpatialGlue’s output. For the Spatial-ATAC-RNA-seq mouse brain dataset, the output obtained from using the same preprocessing steps as SpatialGlue resulted in an excessive number of clusters (more than 20 at a resolution of 0.1). Therefore, we adjusted our preprocessing by first filtering out cells with less than 200 features. Thereafter, we employed SCTransform for normalization and PCA for dimension reduction of the RNA modality. Dimension reduction of the ATAC peaks was performed with Latent Semantic Indexing using the TFIDF and RunSVD functions from the Signac package ^20^. Seurat WNN analysis was then performed with the top 10 PCs of the RNA modality and 2-10PCs of the ATAC modality. We discarded the top ATAC PC as it was correlated with sequencing depth.

## Data availability

All datasets used in this study are already published and were obtained from public data repositories. The SPOTS mouse spleen data was obtained from the GEO repository (accession no. GSE198353, https://www.ncbi.nlm.nih.gov/geo/query/acc.cgi?acc=GSE198353) ^4^, the Stereo-CITE-seq mouse thymus data from BGI and the spatial-ATAC-RNA-seq mouse brain data from AtlasXplore (https://web.atlasxomics.com/visualization/Fan) ^3^. The details of all datasets used are available in the Methods section. The raw data used in this study have been uploaded to Zenodo and is freely available at https://zenodo.org/record/7879713#.ZE3aOnZByUk.

## Code availability

An open-source Python implementation of the GraphST toolkit is accessible at https://github.com/JinmiaoChenLab/SpatialGlue.

## Author contributions

J.C. conceptualized and supervised the project. Y.L. designed the model. Y.L. developed the SpatialGlue software. Y.L., K.S.A., and J.C. wrote the manuscript. Y.L., J.C., R.S, C.Z, H.X., and K.S.A. performed the data analysis. C.Z. and R.S. ran the Seurat WNN algorithm. Y.L. prepared the figures. J.C., N.R.J.G, L.G.N., and N.H. annotated and interpreted the mouse thymus dataset. S.L., Y.H., M.J., A.C., and X.X generated the mouse thymus dataset.

## Acknowledgements

We thank Yingrou Tan for her assistance in mouse thymus data interpretation and Min Wu for his comments on the model. The research was supported by A*STAR under its BMRC Central Research Fund (CRF, UIBR) Award; AI, Analytics and Informatics (AI3) Horizontal Technology Programme Office (HTPO) seed grant (Spatial transcriptomics ST in conjunction with graph neural networks for cell–cell interaction #C211118015) from A*STAR, Singapore; Open Fund Individual Research Grant (Mapping hematopoietic lineages of healthy and high-risk acute myeloid leukemia patients with FLT3-ITD mutations using single-cell omics #OFIRG18nov-0103) from Ministry of Health, Singapore; National Research Foundation (NRF), Award no. NRF-CRP26-2021-0001; the National Research Foundation, Singapore, and Singapore Ministry of Health’s National Medical Research Council under its Open Fund-Large Collaborative Grant (“OF-LCG”) (MOH-OFLCG18May-0003). Singapore National Medical Research Council (#NMRC/OFLCG/003/2018).

## Ethics declarations

The authors declare that there are no competing interests.

## Supplementary Files

Supplementary Figure S1: Results on mouse spleen. (a) Spatial plots of all clusters and each cluster identified by SpatialGlue. (b) DEGs of clusters found in tissue samples. (c) UMAP and spatial plots of clusters identified by Seurat WNN. (d) Modality weights from Seurat WNN explaining the importance of each modality to each cluster.

Supplementary Figure S2: Results on mouse thymus. (a) Spatial plots of all clusters and each cluster identified by SpatialGlue. (b) Expression of marker genes and proteins for each cell type. (c) DEGs of clusters found in tissue samples. (d) UMAP and spatial plots of clusters identified by Seurat WNN. (e) Modality weights from Seurat WNN explaining the importance of each modality to each cluster.

Supplementary Figure S3: Results on mouse brain. (a) Spatial plots of all clusters and each cluster identified by SpatialGlue. (b). DEGs of clusters found in tissue samples. (c) UMAP and spatial plots of clusters identified by Seurat WNN. (d) Modality weights from Seurat WNN explaining the importance of each modality to each cluster.

## Notes

### Competing Interest Statement

The authors have declared no competing interest.

### Summary of Updates

We revised the main manuscript and figures. We added more results.

