## Supplementary figures and images for "Integrated analysis of spatial multi-omics with SpatialGlue"

### Supplementary Figure S1

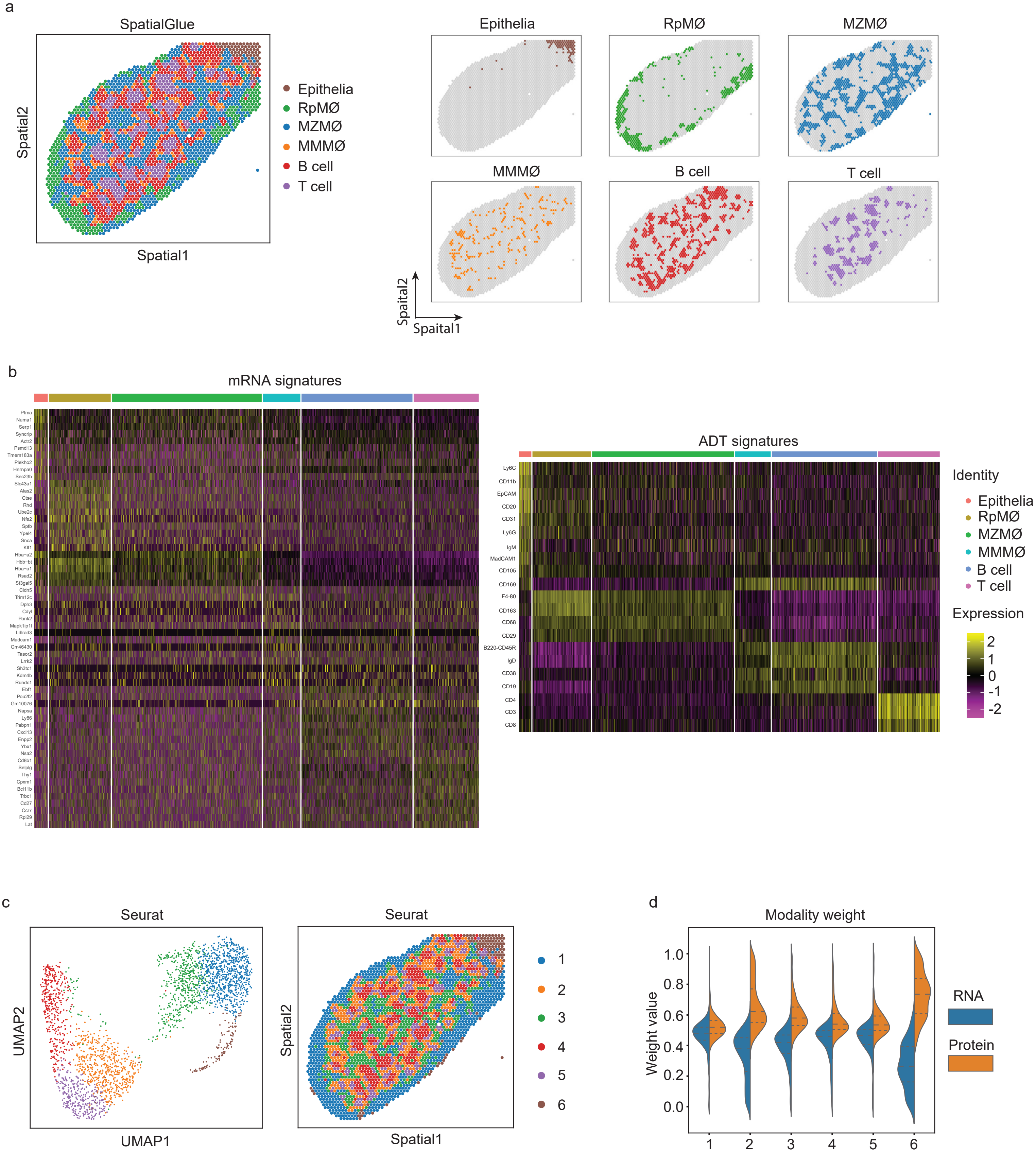

### Supplementary Figure S2

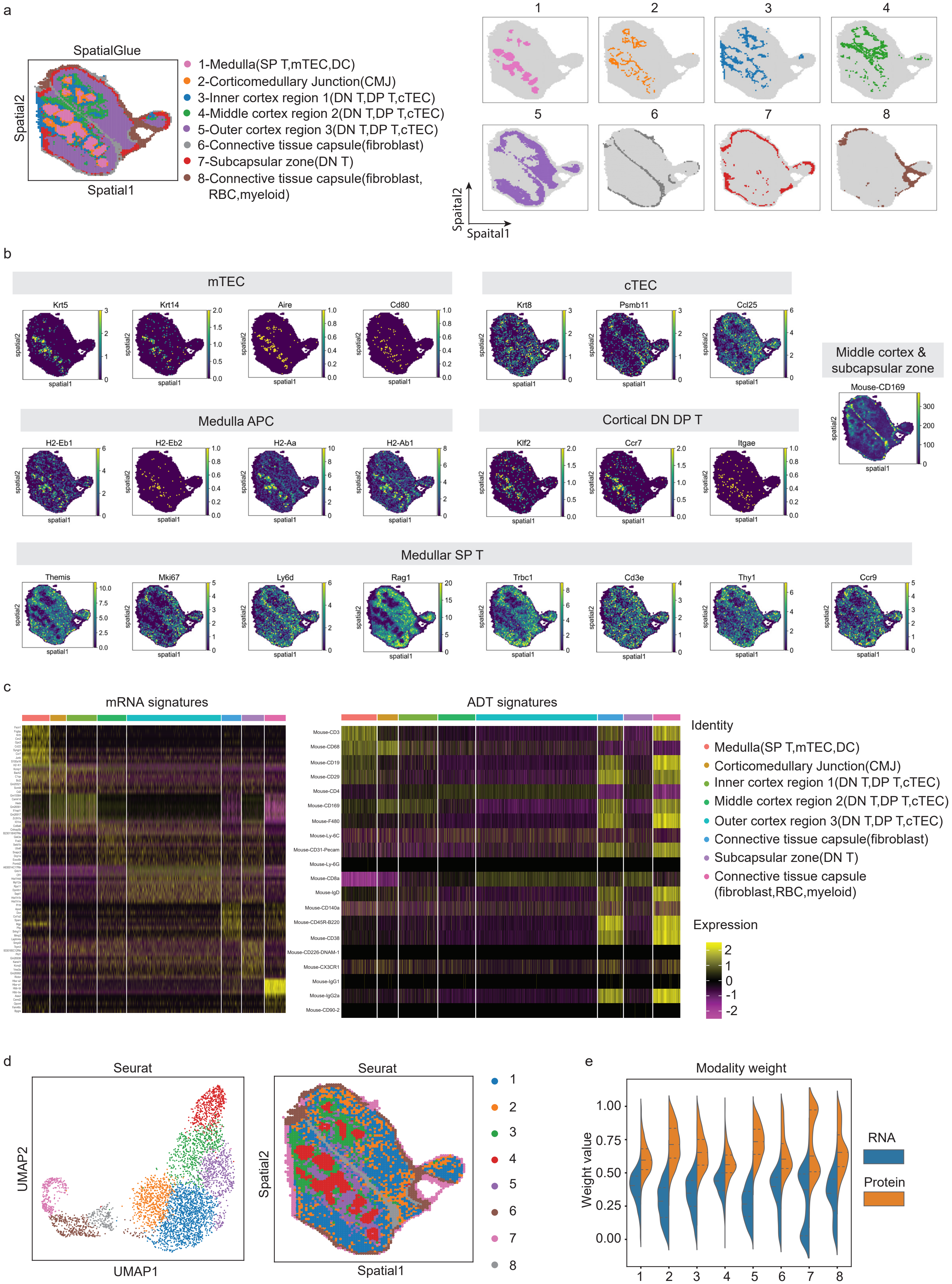

### Supplementary Figure S3

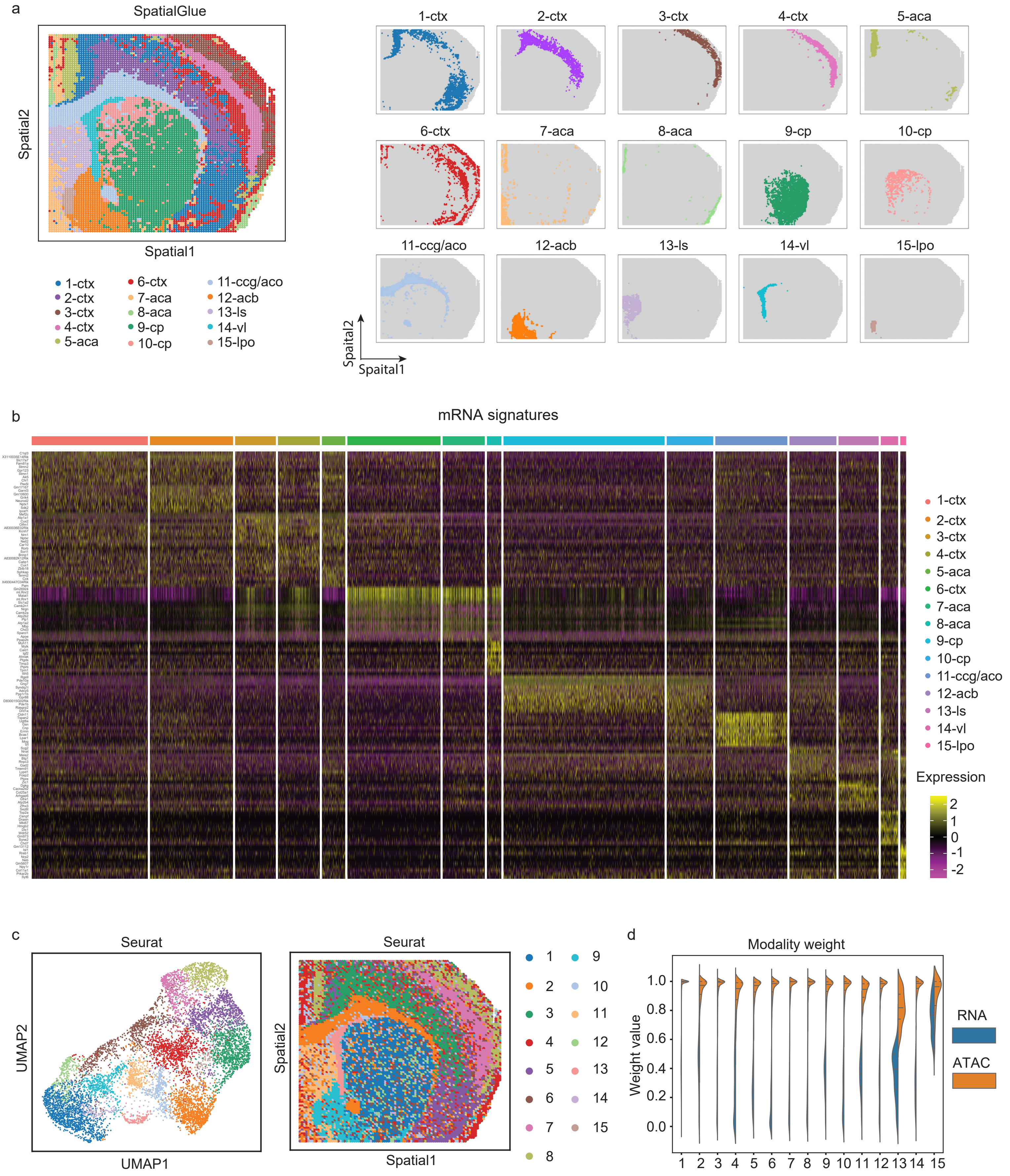
